# T cell-mediated development of stromal fibroblasts with an immune-enhancing chemokine profile

**DOI:** 10.1101/2022.04.03.486884

**Authors:** Ran Yan, Douglas T. Fearon

## Abstract

Stromal fibroblasts reside in inflammatory tissues that are characterized by either immune suppression or immune activation, but whether these cells adapt to these contrasting microenvironments is not known. Cancer-associated fibroblasts (CAFs) mediate immune quiescence by producing the chemokine CXCL12 that coats cancer cells to suppress T cell infiltration. We examined whether CAFs can adopt an immune-promoting chemokine profile. Single-cell RNA-sequencing of CAFs from mouse pancreatic adenocarcinomas identified a sub-population with decreased expression of CXCL12 and increased expression of the T cell-attracting chemokine, CXCL9, that was expanded when tumors were infiltrated with T cells. Conditioned media from activated CD8^+^ T cells containing TNFα and IFNγ converted the chemokine profile of stromal fibroblasts from a CXCL12^+^/CXCL9^-^ immune suppressive phenotype into a CXCL12^-^/CXCL9^+^ immune-activating phenotype. Studies with recombinant cytokines showed that TNFα acted synergistically with IFNγ to induce CXCL9 expression, and alone was mainly responsible for suppressing CXCL12 expression. This coordinated chemokine switch demonstrates that stromal fibroblasts have a phenotypic plasticity that allow their adaptation to contrasting immune tissue microenvironments.

**Summary:** It is unclear whether and how stromal fibroblasts adapt and contribute to varying tissue microenvironments. This study shows immune-suppressive cancer-associated fibroblasts (CAFs) have the capacity to develop an immune-promoting chemokine profile in response to cytokines from infiltrating T cells.

## Introduction

Stromal fibroblasts are present in a range of inflammatory tissues, including cancer, auto-immunity, and infection, that exhibit contrasting immune responses (Buechler et al., 2021; Davidson et al., 2021; Turley et al., 2015). For example, the microenvironment of a tumor is typically immune suppressive (Joyce and Fearon, 2015), whereas that of an infection promotes immune activation. Therefore, a contemporary challenge is to define whether and how the stromal fibroblast adapts and contributes to varying tissue microenvironments. A major step toward answering this challenge was the recognition that subsets of stromal fibroblasts with distinct functions arise in different microenvironments. In a mouse auto-immune arthritis model, two subsets of synovial fibroblasts were identified that mediate joint damage and inflammation, respectively (Croft et al., 2019), and neutralization of TGFβ in a mouse model of breast cancer led to enrichment of a fibroblast subset with a transcriptional profile suggesting signaling by interferon-γ (IFN-γ) (Grauel et al., 2020).

The resistance of many human and mouse carcinomas to inhibition of the PD-1/PD-L1 checkpoint correlates with a relative absence of T cells in cancer cell nests (Fearon, 2017; Joyce and Fearon, 2015). This immune suppressive T cell exclusion phenomenon is mediated, at least in part, by CAF-derived CXCL12 in the form of covalent heterodimers with keratin-19 (KRT19) on the surface of cancer cells (Wang et al., 2022). However, in tissue microenvironments in which T cells accumulate, we hypothesize that CAFs may adopt a contrasting immune-activating phenotype by expressing T cell-attracting chemokines.

## Results and Discussion

We recently reported that a CXCL12/KRT19-coat on mouse pancreatic ductal adenocarcinoma (PDA) cells impairs the accumulation of T cells in nests of cancer cells. CRISPR/Cas9-editing of the *Krt19* gene in mouse PDA cells abolishes formation of the CXCL12/KRT19-coat and allows T cell infiltration in tumors formed with these edited cancer cells (Wang et al., 2022). This circumstance, enabled us to examine the effect of the presence of activated T cells in the tumor microenvironment (TME) on CAFs without perturbing other components of the TME that could alter the CAF phenotype, such as TGFβ (Mariathasan et al., 2018). We formed orthotopic tumors with Scramble-control PDA cells and PDA cells in which *Krt19* had been CRISPR-edited, respectively, and characterized the CAFs from these tumors by single cell RNA-sequencing (scRNA-Seq). Orthotopic tumors formed with Scramble-control PDA cells had cancer cells staining with anti-CXCL12 antibody and few infiltrating T cells. In contrast, in tumors formed with *Krt19*-edited PDA cells, which lacked the CXCL12/KRT19-coat, there were frequent CD3^+^ T cells, and there was decreased tumor growth (Fig. 1A, B; Fig. S1A). Measurement of the mRNA levels of *Cd3e* and *Cd8a* confirmed the immunohistological findings of increased T cells in the immune-activated TMEs, and the increased expression *Ifng* and *Tnf* indicated that these T cells were activated (Fig. 1C). Expression of the two chemokines that have opposing roles in T cell infiltration, *Cxcl9* and *Cxcl12*, were both significantly increased in tumors lacking the CXCL12/KRT19-coat. However, consistent with the accumulation of T cells, the absolute and relative levels of *Cxcl9* (200-fold) were increased more than those of *Cxcl12* (5-fold; Fig. 1C). Therefore, the TMEs of orthotopic tumors formed with Scramble-control PDA cells and *Krt19-*edited PDA cells are immune-quiescent and immune-activated, respectively.

**Figure 1.**
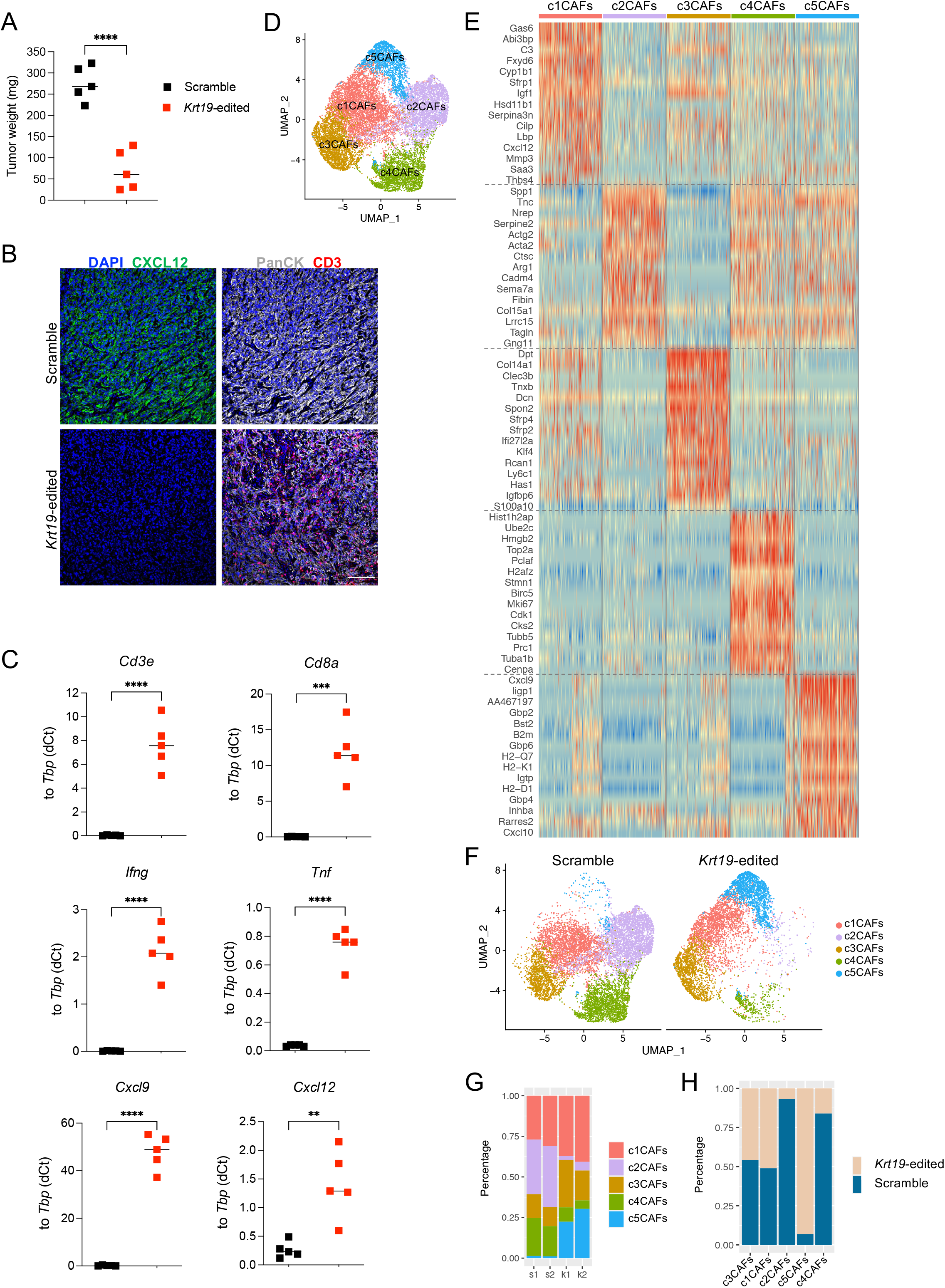
CAFs in T cell-infiltrated, immune-activated, pancreatic tumors have an altered phenotype. **(A, B)** *Krt19* gene editing prevented CXCL12/KRT19-coat formation by cancer cells and reduced PDA growth in the mouse orthotopic tumor model. Unpaired *t*-test analysis on one experiment with five mice per group. Representative images of fluorochrome-conjugated CXCL12, CD3e, and Pan-Cytokeratin antibodies staining on orthotopic tumor sections formed with sgScramble control and sgKRT19-edited PDA cells are shown. Scale bars, 100 μm. **(C)** qPCR analyses of bulk tumors of panel A measuring expression of genes relating to T cells, cytokines, and chemokines are shown. Unpaired *t*-test analysis for one experiment with five mice per group. Figure legend is the same as in (A). **(D)** UMAPs embedding five clusters of 15,738 CAFs from all tumors are shown. Two scramble and five *Krt19*-edited tumors were pooled for one sequencing run. **(E)** The heatmap shows the top 15 enriched genes characterizing each CAF cluster. **(F)** UMAPs embedding the five clusters of 15,738 CAFs separated by the two tumor types, scramble and *Krt19*-edited, are shown. **(G)** Percentage of each CAF cluster in individual sequencing run is shown, wherein each column represents a pooled sequencing sample. **(H)** Percentage of scramble or *Krt19*-edited tumor-derived CAFs within each cluster is shown, wherein each column represents a CAF cluster. ** *p*<0.01, *** *p*<0.001, **** *p*<0.0001.

We measured gene expression of individual cells from these tumors, which were enriched for CAFs by depleting cancer cells and immune cells, and visualized the CAF subpopulations by Uniform Manifold Approximation and Projection (UMAP). All CAF clusters expressed the CAF markers, *Acta2, Fap, Pdpn*, and *Pdgfra*, and did not express *Cdh5, Pecam1, Rgs4* or *Rgs5*, markers of endothelial cells and pericytes (Fig. S1B). When scRNA-seq data from two different runs were integrated, five CAF clusters were apparent (Fig. 1D) with each cluster demonstrating a unique gene expression profile (Fig. 1E). Notably, the relative frequency of CAFs in three of these clusters differed, when the two tumor types were separately assessed; immune-activated tumors contained a higher proportion of c5 CAFs and reduced proportion of c2 and c4 CAFs (Fig. 1F-H).

The gene expression profiles of the five CAF clusters can be related to previously described subpopulations (Biffi et al., 2019; Dominguez et al., 2020; Elyada et al., 2019; Grauel et al., 2020). CAF clusters c1 and c3 resemble iCAFs; the c2 cluster is similar to myCAFs; the c4 cluster is like proliferating CAFs (ProlifCAF); and the c5 cluster resembles interferon-licensed CAFs (ilCAF) (Fig. 2A, B). We termed the latter cluster T-CAF, in recognition of its greater frequency in the T cell-infiltrated tumors. Interestingly, the T-CAFs express CXCL9, a chemokine that would attract CXCR3-expressing T cells (Chow et al., 2019), indicating a positive feedback aspect to CAF differentiation. Since *Cxcl9* is also expressed by immune cells (Groom and Luster, 2011b; Spranger et al., 2017), we sought to estimate the relative contribution of T-CAFs to *Cxcl9* expression in the immune-activated TME by RNA-fluorescence *in situ* hybridization (RNA-FISH) (Fig. 2C). The proportion of *Pdgfra*+ CAFs expressing *Cxcl9* increased from 4% in immune-quiescent tumors to 57% in immune-activated tumors. Further, 34% of all *Cxcl9*+ cells expressing this chemokine were *Pdgfra+* (Fig. 2D). The decreased myCAFs and ProlifCAFs may be secondary to the slower growth of the T cell-infiltrated PDA tumors, and, therefore, a lesser requirement for replicating CAFs.

**Figure 2.**
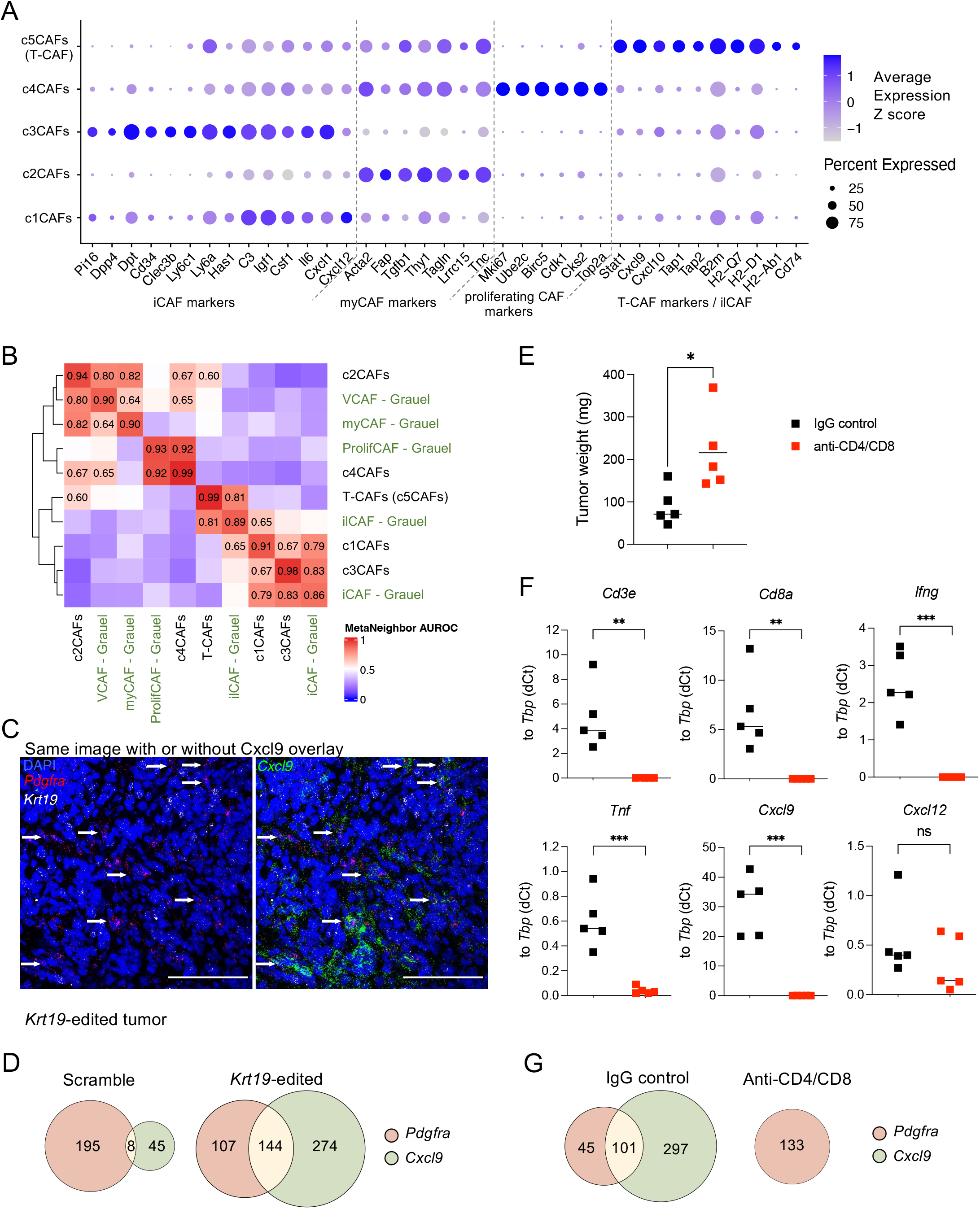
Relationship of the CAF clusters defined by scRNA-seq analyses of immune-quiescent and immune-activated orthotopic PDAs to previously characterized CAF subpopulations. **(A)** The relative expression of iCAF, myCAF, proliferating CAFs and T-CAF marker genes in each CAF cluster are shown. Sizes of the circles represent the percentage of cells expressing the gene in the cluster. Color scale shows the z-score across the row denoting averaging expression level in the cluster. **(B)** Heatmap of area under the receiver operator characteristic curve (AUROC) scores is generated by MetaNeighbor analysis (unsupervised mode) on the most variable genes. 0 indicates no shared identity and 1 indicates identity. Dendrograms were generated by hierarchical clustering of Euclidean distances using average linkage. Row and column labels indicate the gene expression data characterizing previously identified CAF subpopulations. **(C)** Representative images of RNA FISH in a section of an orthotopic *Krt19*-edited PDA in which *Cxcl9* (green), *Pdgfra* (red, indicating CAFs) and *Krt19* (white, indicating PDA cells) mRNA were probed. White arrow indicates *Cxcl9*+ *Pdgfra*+ CAFs. Scale bars, 100 μm. **(D)** Euler diagrams show *Pdgfra*+ only, *Cxcl9+* only or double positive cells in each tumor type. Area of the circle is proportional to cell numbers. Cell numbers were generated by counting 10 images taken from Scramble control *and Krt19*-edited tumors, respectively. **(E)** Weights on day 14 of orthotopically formed, *Krt19*-edited tumors in mice treated with non-immune IgG and anti-CD4/anti-CD8 IgG, respectively, are shown. Unpaired t-test analysis with five mice per group. **(F)** qPCR analyses of tumors of panel E measuring the expression of genes relating to T cells, cytokines, and chemokines are shown. Unpaired *t*-test analysis with five mice per group. **(G)** Euler diagrams of RNA FISH on sections of the tumors from panel E show *Pdgfra*+ only, *Cxcl9+* only, and double positive cells. Area of the circle is proportional to cell numbers.

In T cell-depleted hosts, *Krt19*-edited tumors on day 14 post-implantation were larger and had decreased expression of *Cd3e, Cd8a, Ifng, Tnf* and *Cxcl9* (Fig. 2E-F). By RNA-FISH, *Cxcl9-*expressing CAFs were not detected in these tumors (Fig. 2G), validating the dependence of T-CAFs on intra-tumoral T cells.

To gain insight into how T cells induce T-CAFs, we developed an *in vitro* model in which we modified mouse PDA cells by stably introducing a vector mediating doxycycline-inducible, cytoplasmic ovalbumin (OVA) expression. The induced, OVA-expressing PDA organoids were cultured alone and with effector CD8^+^ T cells expressing an OVA-specific T cell receptor. The conditioned media were collected, and added to mouse pancreatic stellate cells (PSCs), as an example of fibroblastic stromal cells (Fig. 3A) (Ohlund et al., 2017). After six hours, the content in PSCs of *Cxcl9* mRNA was measured. Media from the co-cultures containing antigen-activated CD8^+^ T cells induced *Cxcl9* expression by almost 2000-fold relative to conditioned media from PDA cells alone (Fig. 3B).

**Figure 3.**
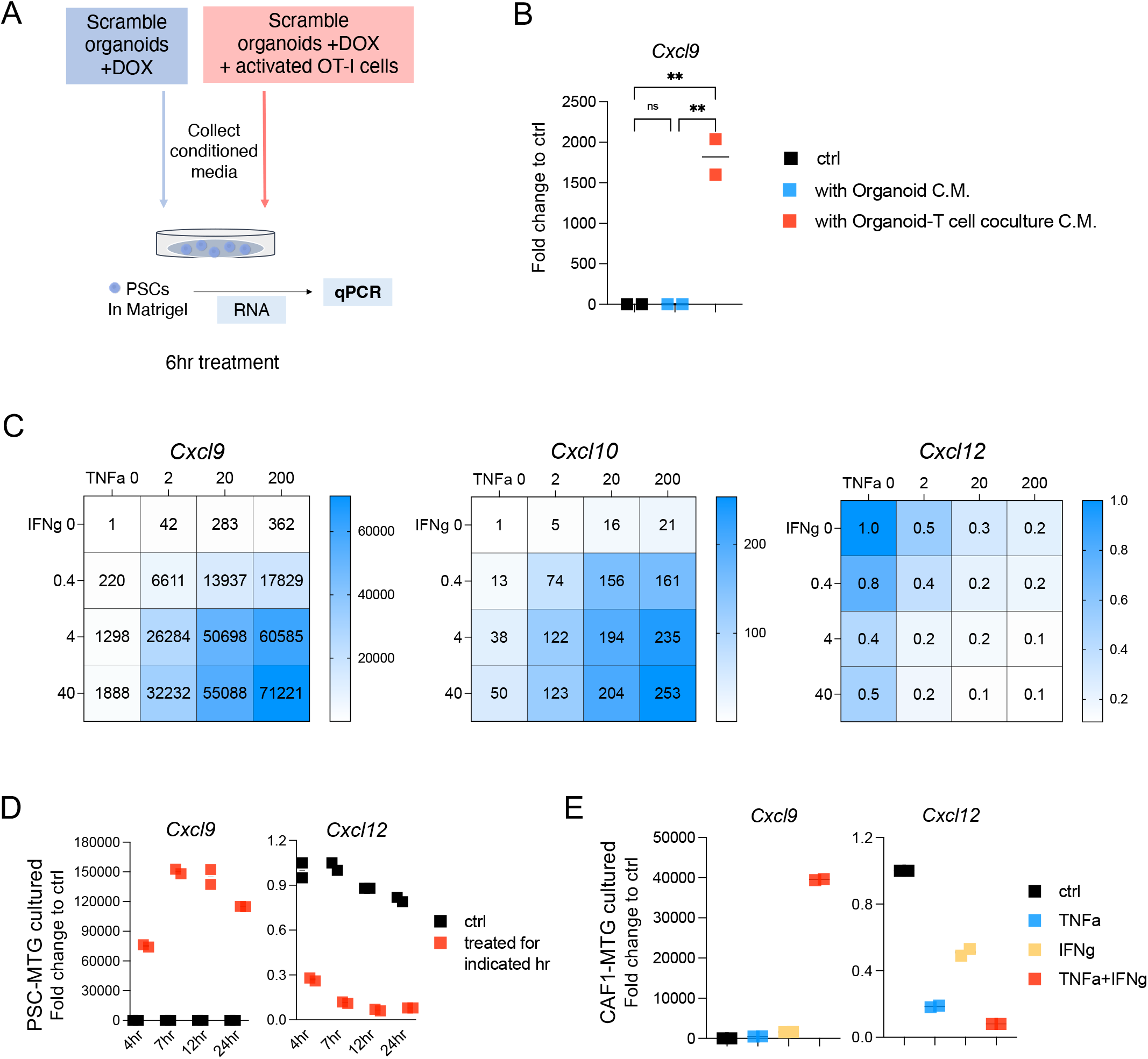
The effects of treating fibroblasts with TNFα and IFNγ on the expression of CXCL9 and CXCL12. **(A)** Schematic is shown of the stimulation of PSCs by conditioned media from cultures of OVA-expressing PDA cells in the absence and presence of *in vitro*-activated OT-I cells. **(B)** qPCR analyses of *Cxcl9* expression in PSCs after treatment with conditioned media. **(C)** Heatmaps show qPCR measurement of *Cxcl9, Cxcl12* and *Cxcl10* expression in Matrigel-cultured PSCs treated with incremental concentrations of TNFα (ng/ml) and IFNγ (ng/ml) alone or together, for 6hrs. Numbers denote fold-change relative to the non-treated controls. **(D)** qPCR analyses of *Cxcl9* and *Cxcl12* expression in Matrigel-cultured PSCs treated with TNFα (4ng/ml) and IFNγ (20ng/ml) for the indicated times. **(E)** qPCR analyses of *Cxcl9* and *Cxcl12* expression in Matrigel-cultured CAF1 cell line treated with TNFα (4ng/ml), IFNγ (20ng/ml), or TNFα and IFNγ in combination for 6hrs.

Two candidates for the cytokines that induce CXCL9 expression by stromal fibroblasts are suggested by three considerations: IFNγ, in combination with TNFα, synergistically induces CXCL9 expression by mice fibroblast cells (Ohmori et al., 1997; Old, 1985); both IFNγ and TNFα mRNA levels are increased in immune-activated PDA tumors (Fig. 1C); and conditioned media from activated CD8^+^ T cell-PDA organoid co-cultures contains increased level of IFNγ and TNFα (Fig. S2A). Therefore, we assessed IFNγ and TNFα, alone and in combination, for their capacity to increase *Cxcl9*. We also assessed their possible effects on the levels of Cxcl12 mRNA and protein expression by PSCs. IFNγ and TNFα synergistically increased CXCL9 expression, whilst TNFα alone was mainly responsible for decreased CXCL12 expression. *Cxcl10* expression was also elevated with the cytokine treatment of PSCs (Fig. 3C, Fig. S2B). The effects of IFNγ and TNFα on CXCL9 and CXCL12 expression in PSCs persisted for up to 24hr (Fig. 3D), and were observed with CAFs in 3-dimensional Matrigel cultures and 2-dimensional plate cultures (Fig. 3E, Fig. S2C-D). TGFβ was unable to block the IFNγ- and TNFα - mediated changes (Fig. S2E).

To determine whether the stromal fibroblasts stimulated by IFNγ and TNFα *in vitro* expressed other genes that characterize T-CAFs, we performed bulk RNA-Seq on PSCs that had been cultured with these two cytokines. Up-regulated mRNAs in the cytokine stimulated PSCs is consistent with the genes defining the T-CAFs (Fig. 4A-B). Also, the observation of a reciprocal relationship of increased *Cxcl9* expression and decreased *Cxcl12* expression by IFNγ- and TNFα -stimulated PSCs and CAFs *in vitro* was observed in UMAPs of CAFs from immune-activated TMEs (Fig. 4C). Therefore, the *Cxcl9*-high, *Cxcl12*-low T-CAF subset may develop in response to IFNγ and TNFα in PDA tumors that are infiltrated with T cells.

**Figure 4.**
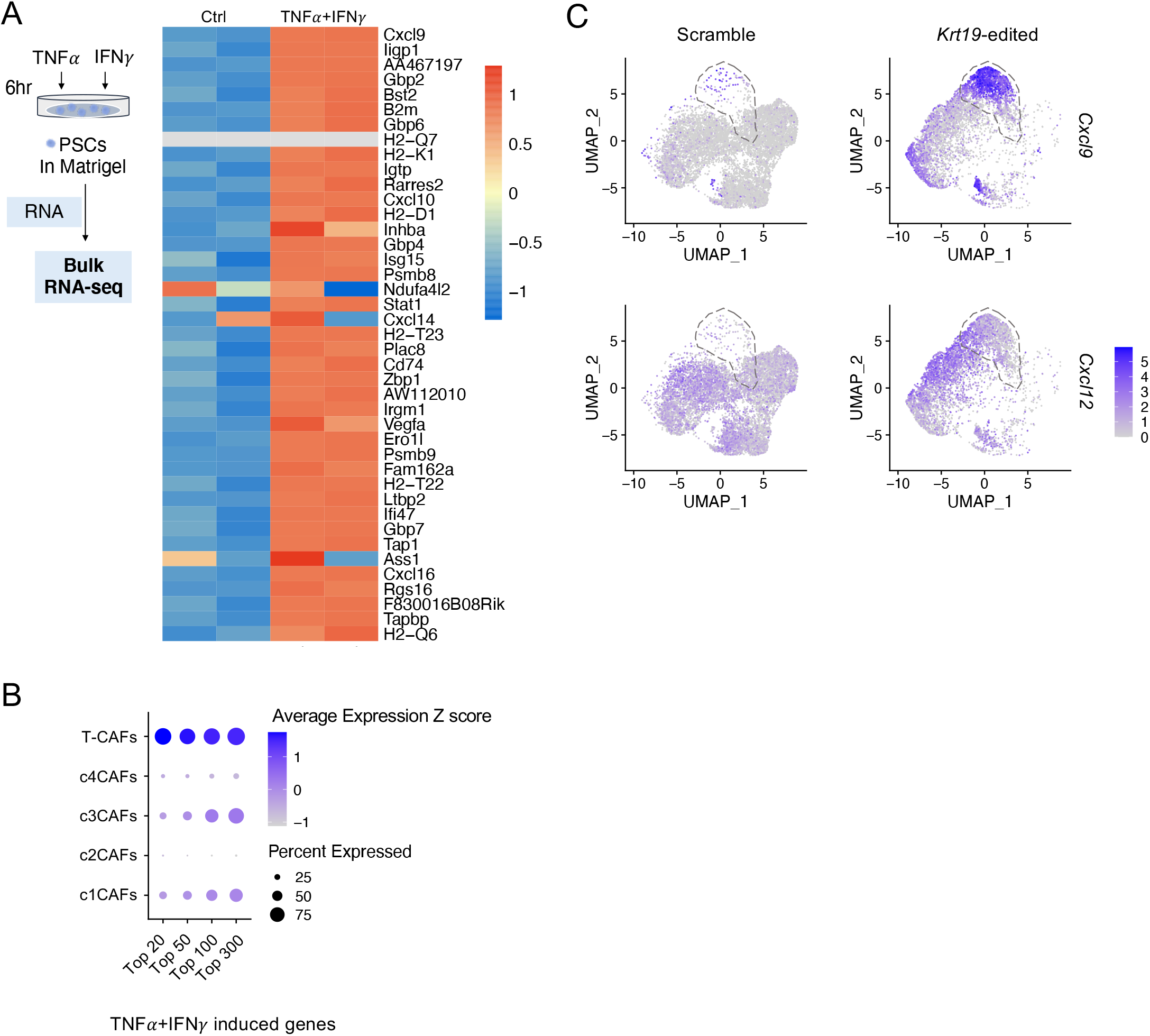
Comparison of fibroblasts stimulated *in vitro* with TNFα and IFNγ with T-CAFs from PDA tumors. **A)** The heatmap shows expression of T-CAF marker gene expression, as revealed by bulk RNA-seq, in TNFα (4ng/ml) and IFNγ (20ng/ml)-treated PSCs. **(B)** The comparison is shown of the mostly highly expressed genes by TNFα - and IFNγ-treated PSCs with the gene signatures in the five CAF clusters. Y axis indicates cluster identity and X axis denotes the number of genes included in the gene signature (20, 50, 100 and 300 top genes). Circle sizes represent percentages of cells expressing the gene signature in a cluster. Color scale shows the z-score across the row denoting averaging expression level in the cluster. **(C)** Expression of *Cxcl9* and *Cxcl12* in CAFs demonstrating reciprocal expression of *Cxcl9* and *Cxcl12* in the c5/T-CAF cluster from *Krt19*-edited PDA tumors.

Orthotopic tumors formed with mouse PDA cells differing only in their expression of *Krt19*, which is required for assembly of the CXCL12/KRT19-coat, had contrasting TMEs that are immune-quiescent and immune-activated (Wang et al., 2022). We use this model to demonstrate two immunological principles. First, stromal fibroblasts have the capacity to sense the presence of activated T cells, independently of other variables, such as TGFβ neutralization (Grauel et al., 2020), and develop a subset that has the potential to promote the T cell reaction. This CAF subset, termed T-CAF, has a chemokine profile that would mediate the accumulation of CXCR3-expressing immune cells, such as NK cells, Th1 cells, CD8^+^ T cells, and dendritic cells (Chow et al., 2019; Groom et al., 2012; Wendel et al., 2008), by coupling the up-regulation of CXCL9 with down-regulation of CXCL12, which otherwise would coat the cancer cells and suppress T cell accumulation (Biasci et al., 2020; Feig et al., 2013; Wang et al., 2022). Interestingly, given its role in auto-immune tissue damage (Feldmann and Maini, 2001; Kalliolias and Ivashkiv, 2016), TNFα is critical in this immunologically cooperative response in that it amplifies the ability of IFNγ to induce the expression of CXCL9 by fibroblasts, including PSCs and CAFs, while suppressing their expression of CXCL12. Thus, the immune-promoting T-CAF develops because these cells integrate signals from two T cell-produced cytokines to present a chemokine profile that could amplify the T cell-mediated immune reaction. Second, this study provides additional support to the concept that the regulation of T cell trafficking is another means by which tissues can determine whether they will be subject to immune-mediated damage (Buckley and McGettrick, 2018; Fearon, 2016; Groom and Luster, 2011a; Peske et al., 2015). The demonstration that the CAF may adopt not only a CXCL12^+^/CXCL9^-^ chemokine profile that protects tumors, as previously shown, but also a CXCL12^-^/CXCL9^+^ phenotype that has the capacity to promote immune attack emphasizes the various roles that stromal fibroblasts may adopt in immune-mediated damage to tissues, including auto-immunity.

## Materials and methods

### Mice

6 week-old male C57BL/6 mice were purchased from Jackson Laboratory. Mice were housed for at least 1 week before each experiment. CD45.1 OT-I mice were generated by cross-breeding OT-I mice (Jackson, 003831) and B6.SJL mice (Jackson, 002014). Protocols for animal experiments were approved by the Institutional Animal Care and Use Committee (IACUC) at Cold Spring Harbor Laboratory (CSHL) in concordance with the NIH Guide for the Care and Use of Laboratory Animals.

### Cell lines and organoids

The clonal PDA cancer cell lines were generated as previously described (Wang et al., 2022). Viral dosages for transduction were reduced 10-fold to obtain low OVA expression lines. Clonal low-OVA lines were generated by treating with doxycycline and selecting single GFP+ cells using flow cytometry. Doxycycline-induced cells were cultured with 5% Tet System Approved FBS (tetracycline-free serum, 631106). To generate tumor organoids, scramble cells were cultured as 3D spheroid in Matrigel for at least 5 passages. OVA and GFP expression was induced by adding 1 ug/ml doxycycline (Takara, 631311) to the culture medium. Pancreatic stellate cell line PSC4 and primary CAF cell line CAF1 were gifts from David Tuveson’s lab. Cells were cultured in DMEM with 5% FBS (Seradigm, 1500-500), 100 units/mL penicillin and 100 μg/mL streptomycin. Cell lines were routinely tested for mycoplasma.

### Mouse tumor generation

For orthotopic tumors, 10,000 2D cultured PDA cells were injected in 20 ul of 50% Matrigel (Corning, 356231) into the pancreas. Tumors were harvested 2 weeks following injection.

### T cell depletion

Mice were given 200μg anti-CD4 (clone GK1.5, BioXcell) and anti-CD8 (clone 53.6-7, BioXcell) antibodies intraperitoneally twice per week, starting two days prior to tumor implantation.

### Immunofluorescence staining

Orthotopic pancreatic tumors were harvested and embedded in Tissue-Tek OCT compound (Sakura, 4583). Fresh frozen tumors were sectioned at 10 μm thickness using a Leica CM3050 cryostat, and sections were stored at -80°C. On the day of staining, sections were incubated with freshly-prepared fixation buffer (4% paraformaldehyde in PBS) for 15 min at room temperature followed by three washes with PBS. Fixed sections were then permeabilized with 0.2% TritonX-100 in PBS for 15 min, followed by blocking with 1% BSA (Calbiochem, 2930) and 10% normal goat serum (Thermo, 16210064) in PBS at room temperature for 1 hour. Sections were incubated with non-conjugated primary CD3 antibody (BioLegend, 100202) at 4°C overnight, and subsequently with fluorophore conjugated secondary antibody (goat anti-rat AF568, Invitrogen A11077) at room temperature for 1 hour. Sections were then incubated with conjugated primary antibodies against CXCL12 (FITC, R&D, IC350F) and pan-Cytokeratin (AF647, Novus, NBP1-48348AF647) at room temperature for 2 hours. Nuclei were stained with DAPI (Thermo, R37606) at room temperature for 10 min. In between each step, sections were washed three times with 0.05% Tween-20 in PBS. Images were acquired on a Leica SP8 confocal microscope using LAS X software at 20x magnification. Images were analyzed with ImageJ software (v2.1.0).

### Tumor digestion and stromal cell enrichment

The tumor digestion protocol was modified from Cremasco *et al*., 2018 (Cremasco et al., 2018). Briefly, tumors were collected, minced, and incubated at 37°C in digestion buffer [RPMI (Gibco), 2% FBS, 0.2 mg/mL Collagenase P (Roche), 0.2 mg/mL Dispase (Gibco), and 0.1 mg/mL DNase I (Roche)] with low agitation (300 rpm) for 45 min, vortexing every 15 min. Released cells were collected from the supernatant and quenched with cold flow cytometry buffer (PBS, 2% FBS, 20 mM HEPES). The remaining tumor pieces underwent a second round of digestion. Released cells from the two round were combined, filtered through 70-um mesh, and concentrated by centrifugation (300 rcf, 5 minutes, 4°C). The single cell suspension was then subjected to ACK lysis and a series of negative selection for removal of dead cells (Miltenyi Dead Cell Removal Kit, cat No. 130-090-101), immune cells (Miltenyi CD45 microbeads, cat No. 130-052-301), and epithelial cells (Miltenyi EPCAM microbeads cat No. 130-105-958). Remaining cells were used for single cell sequencing.

### Single cell RNA sequencing libraries

Single cell RNA-sequencing (scRNA-seq) libraries were prepared from the cell suspension using the Chromium Single Cell 3’ Library V3 kit (10x Genomics). Approximately 10,000 cells per sample were loaded in each lane. Two independent, biological replicates were collected for each condition. Barcoded libraries were pooled and sequenced on NextSeq 2000 P3 flowcell, to a depth of 22,000-56,000 reads per cell.

### RNA FISH

RNA FISH was performed on 10 um fresh frozen tumor sections using the RNAScope Fluorescent Multiplex Reagent kit (ACDBio, 320850). The following mouse RNAScope probes were used: *Cxcl9*-422711, *Pdgfra*-480661-C2, and *Krt19*-402941-C3. Images were acquired on a Leica SP8 confocal microscope at 40x magnification and analyzed with ImageJ.

### Tumor organoid - T cell coculture

OT-I cells were isolated from spleen of an OT-I adult male, using the mouse CD8a+ T Cell Isolation Kit (Miltenyi Biotec, 130-104-075). OT-I cells were than activated using Dynabeads Mouse T-Activator CD3/CD28 (Gibco, 11452D) at a concentration of 1 million/0.5ml in culture media [10% FBS in RPMI with 50uM 2-Mercaptoethanol (Sigma, M6250)] contains 20ng/ml mouse recombinant IL2 (R&D, 402-ML-020). At the same day, about 40,000 scramble tumor organoid cells were seeded in 50ul Matrigel and were cultured in Doxycycline-containing media. Four days later, activated OT-I cells from one well were added to scramble tumor organoids. Conditioned media were collected after 5 days of co-culture.

### Cytokine profiling

Cytokines in conditioned media were detected using the Proteome Profiler Mouse Cytokine Array Kit, Panel A (R&D system, ARY006), following manufacturer’s instructions. X-ray film was scanned on a EPSON scanner (V850 Pro). Integrated pixel density of TNFα and IFNγ spots were measured on inverted images using ImageJ.

### Cytokine treatment and ELISA

10,000 PSC4 cells were seeded in Matrigel one day prior to cytokines treatment. TNFα (R&D, 410-MT-050/CF) and IFNγ (R&D, 485-MI-100/CF) were added to the culture media. CXCL9 concentration was determined using the Mouse CXCL9/MIG DuoSet ELISA kit (R&D, DY492 and DuoSet ELISA Ancillary Reagent Kit 2, DY0081), following manufacturer’s instructions. CXCL12 concentration was determined using the Mouse CXCL12/SDF-1 alpha Quantikine ELISA kit (MCX120, R&D).

### qPCR

RNA extraction was performed using the RNeasy Mini kit (Qiagen, 74106). For tumor samples, ∼30 mg were flash-frozen in liquid nitrogen and stored in -80°C prior to lysis. Cultured cells were directly lysed in RLT buffer (Qiagen). 1 μg total RNA was reverse transcribed with the Taqman Reverse Transcription Kit (Thermo, N8080234), using the oligo d(T) primer. For qPCR, diluted cDNA was mixed with individual Taqman probes (Tbp:Mm00446973_m1, Cd3e:Mm00599683_m1, Cd8a:Mm01182107_g1, Ifng:Mm01168134_m1, Tnf: Mm00443258_m1, Cxcl9: Mm00434946_m1, Cxcl12: Mm00445553_m1) and Taqman Universal Master Mix II (Thermo, 4440040) to a final volume of 10 μL. qPCR reactions were run in 384-well plates using QuantStudio 6. Gene expression was normalized to Tbp.

### Bulk RNA-Seq

Two biological replicates were collected per condition. Sequencing libraries were prepared using the Illumina TruSeq Stranded Total RNA Library Prep kit (20020594). Briefly, 4 μg total RNA per sample was used to prepare cDNA libraries. cDNA was amplified with 12 PCR cycles to minimize duplicate reads. Libraries were pooled and sequenced with single-end 76 bp reads on an Illumina NextSeq 500, using the High Output kit.

### Single cell RNA-seq data processing and analysis

FASTQ files were processed using the Cell Ranger pipeline (v6.0.0) with the mm10-2020-A reference. Ambient RNA was removed from the filtered H5 barcode file using Cellbender remove-background. The expected number of cells and empty droplets were estimated from the UMI curve generated by the Cell Ranger pipeline. Data analysis was performed in R (v4.1.0) using Seurat (v 4.0.5) (Hao et al., 2021). HDF5 files containing count matrices for each sample were loaded into individual Seurat objects and then merged. In total, 28,318 cells passed QC filtering (>200 genes/cell, <10% mitochondrial genes, doublet removal using DoubletFinder v3). Gene expression was normalized and log-transformed, and the top 2000 variable genes were identified using FindVariableFeatures. Gene counts were scaled, and linear dimensionality reduction was performed with principal component analysis (RunPCA). The number of PCs to include for downstream clustering was estimated using the Elbow plot method. To cluster the cells, a KNN graph based on the Euclidean distance in PCA space was generated (FindNeighbors), and clustering was performed with the Louvain algorithm (FindClusters, resolution=0.5). Marker genes for each cluster were calculated using FindAllMarkers function and filtered for genes expressed in ≥25% of the population with ≥log(0.25)-fold higher expression. Marker genes for each CAF population were identified using FindMarkers function, comparing each CAF cluster to the remaining CAF clusters and again filtering for genes expressed in ≥25% of the population with ≥log(0.25)-fold higher expression. Expression of the TNFα /IFNγ signature was calculated per cell using the AddModuleScore function. MetaNeighbor (v1.10.0) (Crow et al., 2018) was used to compare CAF clusters to a published CAF dataset (Grauel et al., 2020 [GSE160687]). Briefly, the two datasets were merged into one SingleCellExperiment object. Unsupervised MetaNeighbor analysis was performed between CAF clusters identified here and in Grauel et al. 2020, using highly variable genes identified across datasets (called with ‘variableGenes’ function). The heatmap of MetaNeighbor AUROC values was generated with ComplexHeatmap (Gu et al., 2016).

### Bulk RNA-seq data processing and analysis

FASTQ files were loaded into CSHL Galaxy workspace (v1.0.0). After pass QC (FastQC), reads from each sample were aligned to the GRCm38/mm10 reference genome using STAR (Galaxy Tool Version: 2.5.2b-1)(Filter by number of mismatches=2; Maximum number of alignments allowed=2). Read counts over gene exons were detected using featureCounts (version 1.4.6.p5). Differential gene expression was performed using DESeq2 (Galaxy Tool ID: cshl_DESeq2_1).

### Statistical analysis

Prism was used for performing statistical analysis and graphical representation of data. Student t test and one-way ANOVA were used depending on the sample size.

## Online supplemental material

Fig. S1 The altered phenotype of CAFs in T cell-infiltrated Krt19-edited tumors.

Fig. S2 TNFα and IFNγ on CXCL9 and CXCL12 expression in fibroblasts *in vitro*.

## Acknowledgments

We thank Bruno Gegenhuber for help with scRNA-seq analysis; Dr. David Tuveson for providing cell lines and valuable suggestions; members in the Fearon lab for insightful discussion; Dr. Hannah Meyer for reviewing the manuscript. This work was performed with assistance from CSHL Animal, Histology, Microscopy, Flow Cytometry, Single Cell Biology, and the Sequencing Technologies and Analysis shared resources, which are supported by the Cancer Center Support Grant 5P30CA045508.

## Funding

Distinguished Scholar Award of the Lustgarten Foundation (DTF)

Thompson Family Foundation (DTF)

National Cancer Institute (DTF)

George A. & Marjorie H. Anderson Scholarship from the School of Biological Sciences (RY)

