## supplemental figure legend for "T cell-mediated development of stromal fibroblasts with an immune-enhancing chemokine profile"

Figure S1.

**The altered phenotype of CAFs in T cell-infiltrated *Krt19*-edited tumors.**

**(A)** Images are shown of orthotopic scramble-control tumors and *Krt19*-edited tumors generated for scRNA-seq analyses, with weights of individual tumors. Unpaired *t* test analysis on the weight of 10 tumors from two experiments combined. Two scramble-control tumors (marked with red asterisk) and five *Krt19*-edited tumors, respectively, were pooled for one sequencing run. **(B)** Expression of marker genes for different cell types from the two tumor types is presented in a Dotplot. The sizes of circles represent percentages of cells expressing the gene in a cluster. Color scale shows the z-score across the row denoting average expression level in the cluster. **** *p*<0.0001

Figure S2.

**TNFα and IFNγ on CXCL9 and CXCL12 expression in fibroblasts *in vitro.***

**(A)** Quantification of TNFα and IFNγ dot pixel density from cytokine array analysis on collected conditional media (same in Fig 3B) demonstrates increased TNFα and IFNγ protein in conditioned media from PDA organoid/OT-I cell co-cultures. **(B)** Heatmaps show amounts of secreted CXCL9 and CXCL12 in media from Matrigel-cultured PSCs treated with different concentrations of TNFα (ng/ml) and IFNγ (ng/ml) for 6 hrs. Numbers denote detected chemokine concentration (ng/ml). **(C)** qPCR analyses are shown of *Cxcl9* and *Cxcl12* expression in 2D-cultured PSCs treated with TNFα (4ng/ml), IFNγ (20ng/ml), and TNFα and IFNγ in combination. **(D)** qPCR analyses are shown of *Cxcl9* and *Cxcl12* expression in 2D-cultured CAF1 cell line treated with TNFα (4ng/ml), IFNγ (20ng/ml), and TNFα and IFNγ in combination. **(E)** qPCR analyses are shown of *Cxcl9* and *Cxcl12* expression in Matrigel-cultured PSCs treated with TNFα (4ng/ml) and IFNγ (20ng/ml) in the presence or absence of TGFβ (50ng/ml) for 6hrs.
