## Supplementary figures and images for "T cell-mediated development of stromal fibroblasts with an immune-enhancing chemokine profile"

### supplemental figure 1 and 2

#### Supplemental figure 1

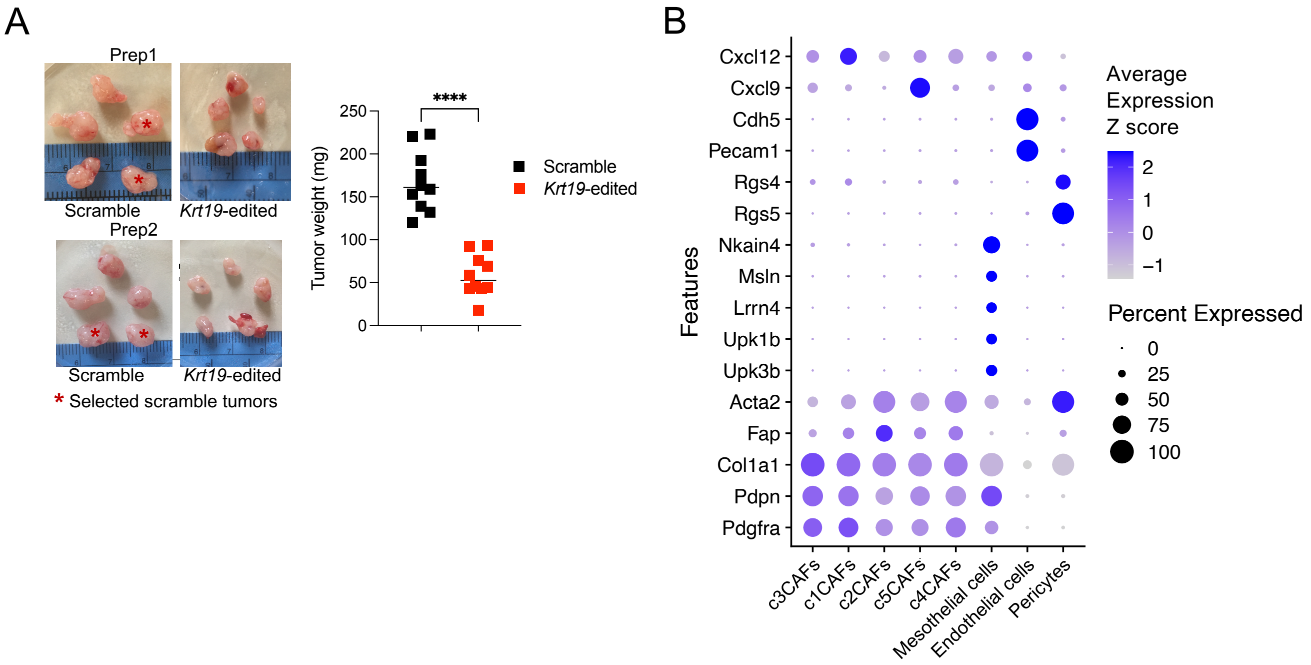

#### Supplemental figure 2

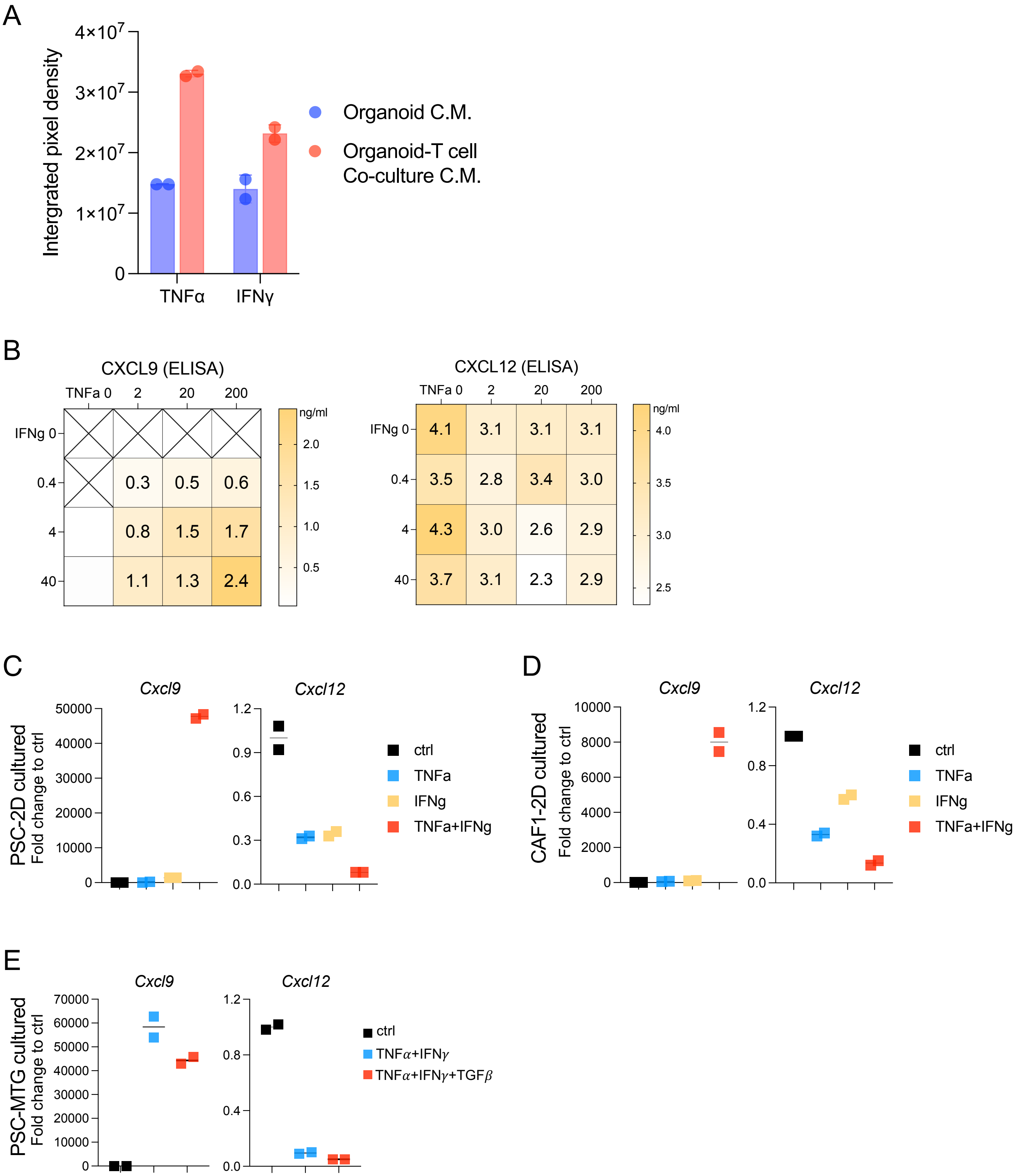
